# Refactoring gene sequences for broad assembly standards compatibility

**DOI:** 10.1101/225284

**Authors:** Tyson R. Shepherd

**Affiliations:** Department of Biological Engineering, Massachusetts Institute of Technology, Cambridge, MA 02139, USA

## Abstract

Four cloning standards in synthetic biology are BioBrick, BglBrick, MoClo and GoldenBraid, with each requiring their constitutive parts be compatible with the associated restriction enzymes. To standardize parts for the broadest usage, it would be useful to synthesize genes that are simultaneously compatible with all 4 popular assembly strategies. Here it is shown that using a defined set of rules, implemented in a computational program, any protein coding sequence can be made compatible with all four standards by silent mutations. Using a coding sequence as an input, all BioBrick, BglBrick, MoClo, and GoldenBraid restriction sites and chi recombination hot spots can be destroyed with silent mutations that approximate the codon usage of the organism. As an application, all open reading frames in the model organisms *Escherichia Coli* and *Bacillus Subtilis* are computationally refactored, showing the feasibility of implementing this umbrella strategy for synthesizing genes with the broadest compatibility.

## Introduction

One of the many goals of synthetic biology is the standardization of gene sequences encoding characterized biochemical parts to ease distribution and standardize assemblies^1–2^. Traditional assembly standards in the community include BioBrick, BglBrick, and type IIs-based cloning (e.g., MoClo and GoldenBraid), each having notable advantages. BioBrick cloning uses classic restriction cloning for idempotent assemblies, showing utility for synthesizing systems up to 30 cistrons^3–6^, and already has wide acceptance with a community parts resource (http://parts.igem.com). BglBricks also uses standard idempotent cloning that offers in-frame assembly of parts^7^. Type IIs methods such as MoClo and GoldenBraid assemblies offer a single-pot cloning of larger systems, allowing for combinatorial cloning of many parts simultaneously^8–11^.

The lowering cost and wider availability of DNA synthesis has allowed for the democratization of gene and pathway synthesis, which has been met with development of several useful computer aided design synthetic biology tools^12–14^. However, this has increased the need for standards to improve reproducibility while reducing redundant synthesis of parts. This is especially important in the large-scale synthesis of parts^10, 15^ and in refactoring synthetic genomes^5, 16–19^. However, the compatibility of the four widely-used standards to be simultaneously implemented in a single dictionary of gene sequences has not been reported. Here, a computational algorithm (Synthetic biology Gene Standardizer or “SyGS”) following a defined ruleset is used to make silent mutations at every BioBrick, BglBrick, MoClo, and GoldenBraid restriction site for any given protein coding sequence. The algorithm additionally allow removal of all crossover hotspot instigator (chi) recombination sites^20^ and, in *Escherichia coli (E. coli)*, any annotated internal transcription promoters^21^. To test the feasibility of applying this umbrella strategy generally, every open reading frame (ORF) of two widely used model organisms, *E. coli* and *Bacillus subtilis* (*B. subtilis*), are processed by the program, removing all targeted sites without any necessary amino acid substitutions. The results are part of a reference dictionary hosted at http://trsheph.scripts.mit.edu/SyGS/.

## Results and Discussion

SyGS v1.0 is a computational sequence-refactoring tool that automates the standardization of gene sequences to have the broadest compatibility with current synthetic biology cloning strategies (**Fig 1**). The program searches for restriction sites used in BioBrick (*Eco*RI, *Xba*I, *Spe*I, *Pst*I, *Apo*I, *Mfe*I, *Avr*II, *Nhe*I, *Nsi*I, *Sbf*I, *Not*I), BglBrick (*Eco*RI, *Bgl*II, *Bam*HI, *Xho*I), MoClo (*Bsa*I, *Bbs*I, *Mly*I), and GoldenBraid (*Bsa*I, *Bsm*BI, *Btg*ZI) cloning. Additionally, all *E. coli* chi sites are removed to reduce recombination ^20^ and enable further engineering^22^. All start codons are unified to ATG and all stop codons are set to TAA. *Nde*I sites are also removed for ease of assembly analysis and cloning at the ATG start codon. Once identified, silent mutations to a codon within each targeted site are made to an alternative codon with its choice based on removing the target site and matching the statistics of the codon usage of the organism [http://www.genscript.com/tools/codon-frequency-table], although a minimal algorithm can be used to remove all sites by changing one codon to its most frequently used codon (**Table 1**).

**Figure 1.**
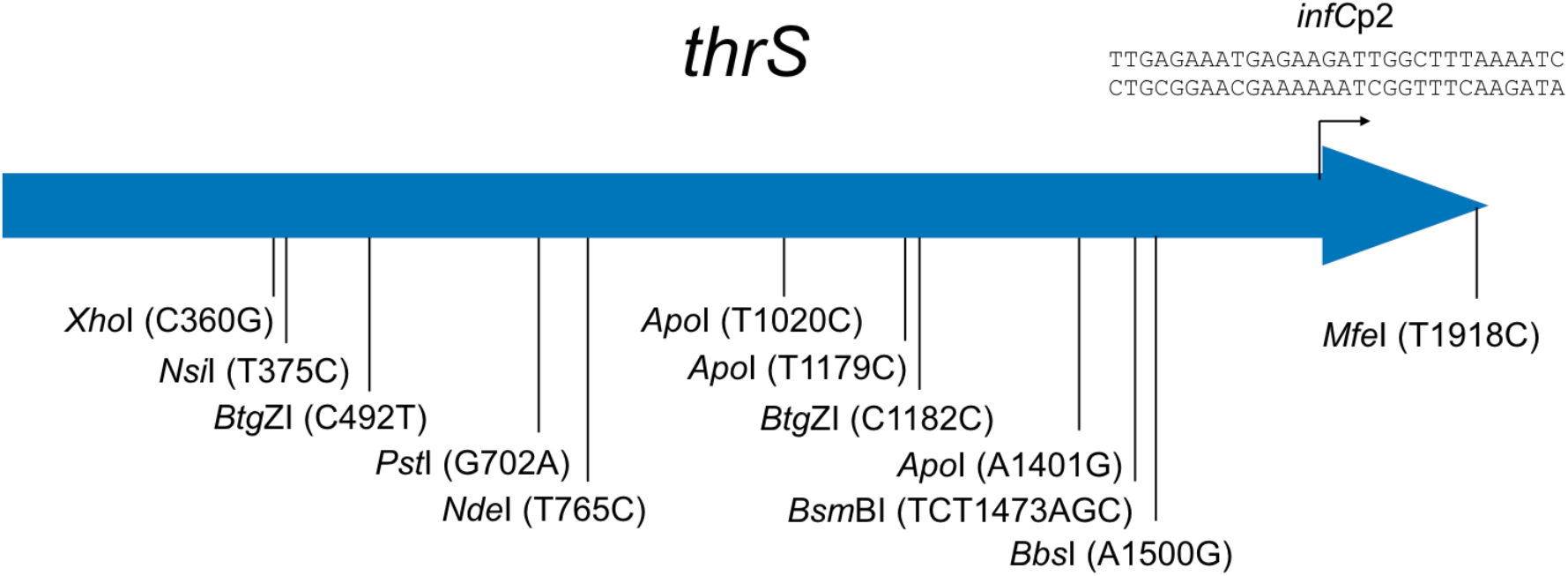
Example of mutations to refactor *E. coli thrS* (threonine-tRNA ligase). Twelve restriction sites are eliminated by silent mutations to the cut sites as labeled. Annotated internal promoters in *E. coli* genes, such as the shown *inf*Cp2, are also silently disrupted.

**Table 1.**
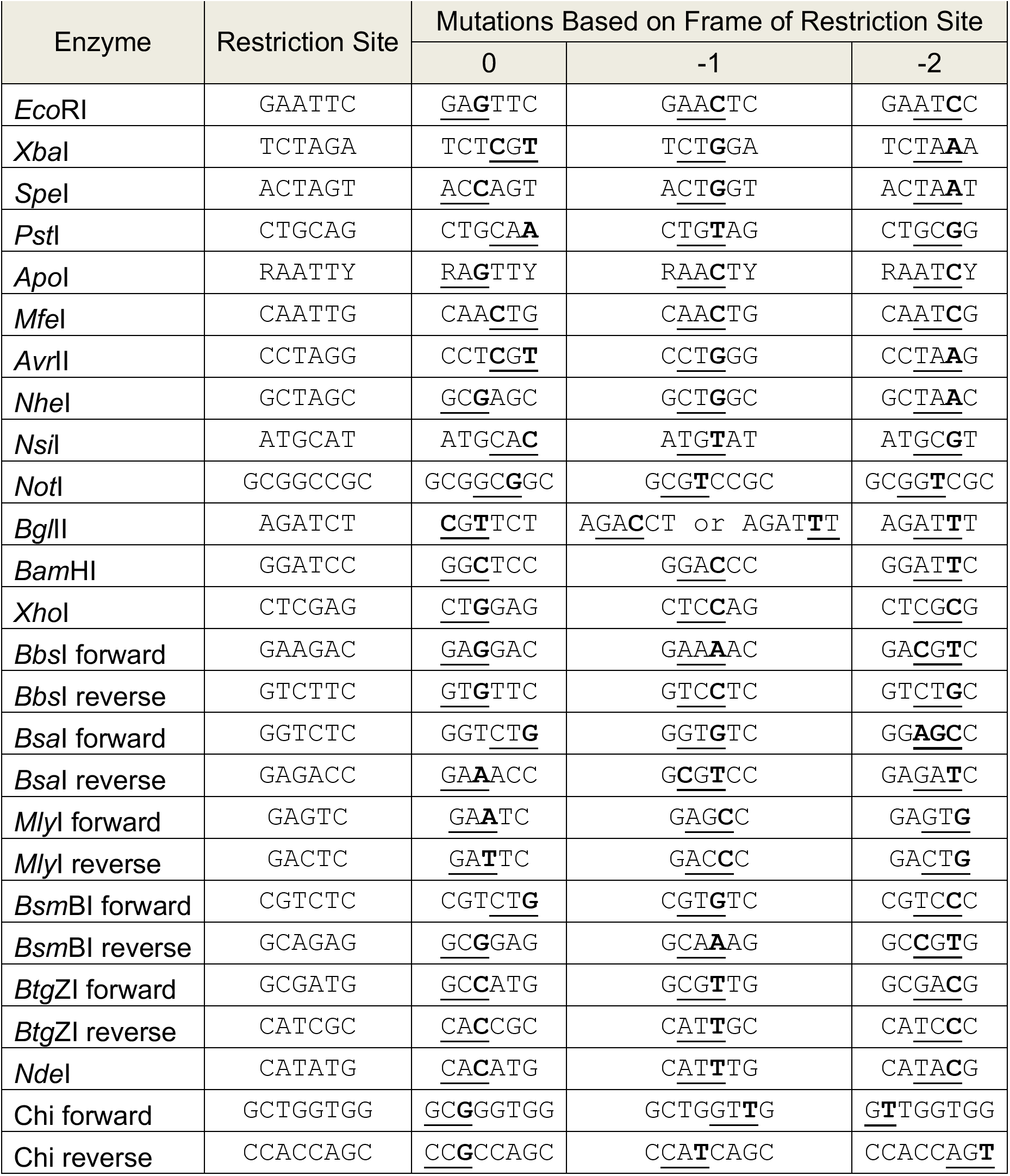
Minimal rule set to automate mutations for gene synthesis. Underline indicates the codon in the ORF targeted for mutation. Mutations are in bold.

Next, the entire ORF libraries of *E. coli* and *B. subtilis* were processed by the program. The output was the refactoring of every ORF sequence to be compatible with four current synthetic biology assembly standards. In all, 22,748 mutations are suggested for *E. coli* (about 5 per ORF on average), while 24,397 mutations are suggested for *B. subtilis* (**Fig. 2**).

**Figure 2.**
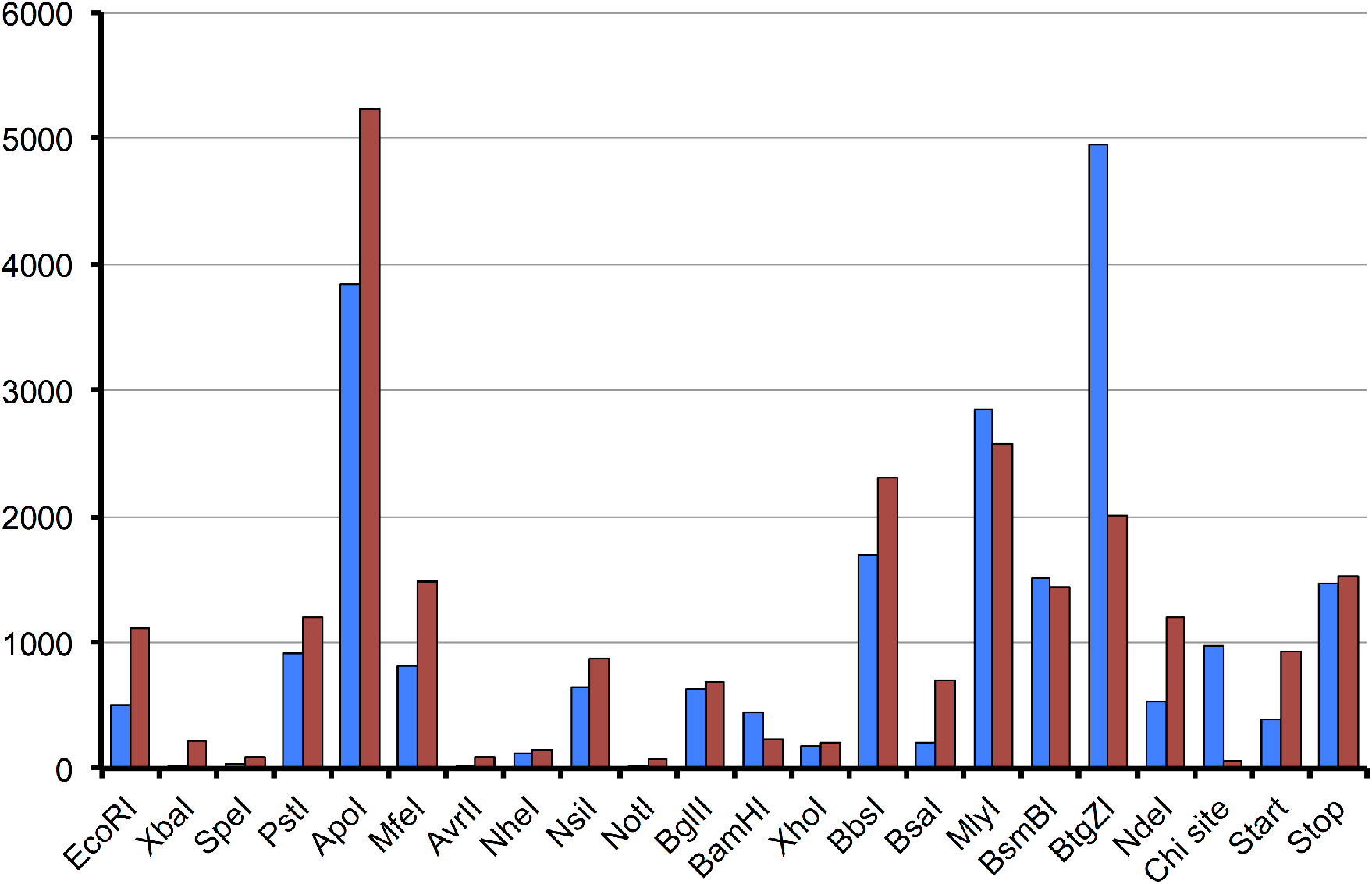
Occurrence of suggested mutations per restriction site to refactor all ORFs in *E. coli* (Blue) or *B. subtilis* (Red) to be compatible with four major synthetic biology assembly standards as well as removing NdeI, chi sites, and unifying start and stop codons.

Internal transcription initiation sites are also problematic in systems assembly of parts^5^. To mitigate this problem, the sequences of *E. coli* ORFs were cross-checked against the RegulonDB promoter database^21, 23^, and any annotated internal promoter sequences were silently mutated to remove them.

Access to any of the refactored *E. coli* and *B. Subtilis* ORF sequences and to refactor any new sequence is offered as a web resource at http://trsheph.scripts.mit.edu/SyGS/. This online resource allows a user to enter a gene name for either *E. coli* or *B. subtilis*, or paste a sequence into a form that, once submitted, returns the synthetic sequence that has been refactored to be free from all sites listed above. Additionally, the output shows the suggested mutations for each of the sites that were modified. The command line source code for batch processing is available at https://github.com/trsheph/SynBioStandardizer.

As codon optimization of the remainder of the sequence is not taken into consideration here, it is suggested that sequences from alternative organisms may benefit by first running the sequence through a codon optimizer.

As gene synthesis becomes more prevalent, care should be taken to generate genes with the broadest compatibility with current assembly standards. SyGS v1.0 makes the task of refactoring of any sequence of interest to be compatible with four major synthetic biology assembly standards quick and easy.

## Materials and Methods

### Program method

SyGS is written in the python programming language and uses the BioPython library^24^ for searching for restriction sites and manipulating the underlying sequence. The command line program input is a text file containing one or more fasta formatted sequences. This data is parsed into a SeqIO record object and the sequence is searched for restriction sites used in BioBrick, BglBrick, MoClo, and GoldenBraid assemblies (**Table 1**). Additionally, the sequence is searched for *Nde*I sites and *E. coli* chi sites (5’-GCTGGTGG-3’). The placement of every target site within the reading frame is determined and one codon in the site is modified to a different synonymous codon selected based on the statistics of the codon usage of *E. coli*. A special case exists for BsaI and *Bgl*II, where the sequence 5’-GAGATCTN-3’ in the -1 frame will only allow a silent leucine codon mutation from CTN to a split codon TTG, the G being outside of the restriction sequences. This takes the more frequently used leucine codons to a 10% usage codon. No conflicts exist between the other targeted sequences using the minimal rule set in **Table 1**.

The refactoring is then crosschecked by comparing the protein translation of the refactored sequence to the translation of the input sequence. The program output on the command line is “successful” if the translations are the same and there are no sequence conflicts during mutations. The output is a text file containing the refactored sequences, a second information text file containing the suggested mutations, and a third statistics file.

### Refactoring all ORFs of an organism

Complete coding sequences records for *E. coli* str. K-12 substr. MG1655 (NC_000913.3) and *B. subtilis* str. 168 (NC_000964.3) were downloaded from NCBI as text files. These were each processed via the command line interface, with the program processing some 4,000 ORFs in less than one minute. *E. coli* sequences were additionally cross-checked against the internal transcription promoter database^23^ and mutations were made to remove any internal promoters. The output was a text file containing fasta-formatted synthetic sequences for each of the ORFs in the downloaded file and a second file containing each suggested mutation.

### Online resource

*E. coli* or *B. subtilis* refactored sequences and mutations can be displayed by selecting the organism and the name of the gene. Alternatively, the user can submit any coding sequence in a text box, which the script will operate on and return the new suggested refactored sequence and the accompanying mutations.

## Acknowledgements

I am grateful for the support from Prof. Anthony Forster (Uppsala University, Uppsala, SE) and Prof. Mark Bathe (Massachusetts Institute of Technology, MA, USA) during development this work, and the support by a Wenner-Gren Foundation postdoctoral fellowship to T.S. and to ONR (grants N00014-16-1-2953 and N00014-17-1-2609) to Prof. Mark Bathe.

## Source code availability

The source code for SyGS is open source under the BSDv2 license and can be found in GitHub at https://github.com/trsheph/SynBioStandardizer

## References

1. Arkin, A., Setting the standard in synthetic biology. Nature biotechnology 2008, 26 (7), 771–4.

2. Purnick, P. E.; Weiss, R., The second wave of synthetic biology: from modules to systems. Nature reviews. Molecular cell biology 2009, 10 (6), 410–22.

3. Knight, T., Idempotent vector design for standard assembly of biobricks. 2003.

4. Shetty, R. P.; Endy, D.; Knight, T. F., Jr., Engineering BioBrick vectors from BioBrick parts. Journal of biological engineering 2008, 2, 5.

5. Shepherd, T. R.; Du, L.; Liljeruhm, J.; Samudyata; Wang, J.; Sjodin, M. O. D.; Wetterhall, M.; Yomo, T.; Forster, A. C., De novo design and synthesis of a 30-cistron translation-factor module. Nucleic Acids Res 2017, 45 (18), 10895–10905.

6. Popp, P. F.; Dotzler, M.; Radeck, J.; Bartels, J.; Mascher, T., The Bacillus BioBrick Box 2.0: expanding the genetic toolbox for the standardized work with Bacillus subtilis. Sci Rep 2017, 7 (1), 15058.

7. Anderson, J. C.; Dueber, J. E.; Leguia, M.; Wu, G. C.; Goler, J. A.; Arkin, A. P.; Keasling, J. D., BglBricks: A flexible standard for biological part assembly. Journal of biological engineering 2010, 4 (1), 1.

8. Engler, C.; Kandzia, R.; Marillonnet, S., A one pot, one step, precision cloning method with high throughput capability. PloS one 2008, 3 (11), e3647.

9. Sarrion-Perdigones, A.; Falconi, E. E.; Zandalinas, S. I.; Juarez, P.; Fernandez-del-Carmen, A.; Granell, A.; Orzaez, D., GoldenBraid: an iterative cloning system for standardized assembly of reusable genetic modules. PloS one 2011, 6 (7), e21622.

10. Smanski, M. J.; Bhatia, S.; Zhao, D.; Park, Y.; L, B. A. W.; Giannoukos, G.; Ciulla, D.; Busby, M.; Calderon, J.; Nicol, R.; Gordon, D. B.; Densmore, D.; Voigt, C. A., Functional optimization of gene clusters by combinatorial design and assembly. Nature biotechnology 2014, 32 (12), 1241–9.

11. Weber, E.; Engler, C.; Gruetzner, R.; Werner, S.; Marillonnet, S., A modular cloning system for standardized assembly of multigene constructs. PloS one 2011, 6 (2), e16765.

12. Cai, Y.; Wilson, M. L.; Peccoud, J., GenoCAD for iGEM: a grammatical approach to the design of standard-compliant constructs. Nucleic Acids Res 2010, 38 (8), 2637–44.

13. Hillson, N. J.; Rosengarten, R. D.; Keasling, J. D., j5 DNA assembly design automation software. ACS synthetic biology 2012, 1 (1), 14–21.

14. Xia, B.; Bhatia, S.; Bubenheim, B.; Dadgar, M.; Densmore, D.; Anderson, J. C., Developer’s and user’s guide to Clotho v2.0 A software platform for the creation of synthetic biological systems. Methods Enzymol 2011, 498, 97–135.

15. Temme, K.; Zhao, D.; Voigt, C. A., Refactoring the nitrogen fixation gene cluster from Klebsiella oxytoca. Proc Natl Acad Sci U S A 2012, 109 (18), 7085–90.

16. Forster, A. C.; Church, G. M., Towards synthesis of a minimal cell. Mol Syst Biol 2006, 2, 45.

17. Forster, A. C.; Church, G. M., Synthetic biology projects in vitro. Genome Res 2007, 17 (1), 1–6.

18. Jewett, M. C.; Forster, A. C., Update on designing and building minimal cells. Curr Opin Biotechnol 2010, 21 (5), 697–703.

19. Ostrov, N.; Landon, M.; Guell, M.; Kuznetsov, G.; Teramoto, J.; Cervantes, N.; Zhou, M.; Singh, K.; Napolitano, M. G.; Moosburner, M.; Shrock, E.; Pruitt, B. W.; Conway, N.; Goodman, D. B.; Gardner, C. L.; Tyree, G.; Gonzales, A.; Wanner, B. L.; Norville, J. E.; Lajoie, M. J.; Church, G. M., Design, synthesis, and testing toward a 57-codon genome. Science 2016, 353 (6301), 819–22.

20. Dillingham, M. S.; Kowalczykowski, S. C., RecBCD enzyme and the repair of double-stranded DNA breaks. Microbiol Mol Biol Rev 2008, 72 (4), 642–71, Table of Contents.

21. Huerta, A. M.; Collado-Vides, J., Sigma70 promoters in Escherichia coli: specific transcription in dense regions of overlapping promoter-like signals. J Mol Biol 2003, 333 (2), 261–78.

22. Marshall, R.; Maxwell, C. S.; Collins, S. P.; Beisel, C. L.; Noireaux, V., Short DNA containing chi sites enhances DNA stability and gene expression in E. coli cell-free transcription-translation systems. Biotechnology and bioengineering 2017, 114 (9), 2137–2141.

23. Gama-Castro, S.; Jimenez-Jacinto, V.; Peralta-Gil, M.; Santos-Zavaleta, A.; Penaloza-Spinola, M. I.; Contreras-Moreira, B.; Segura-Salazar, J.; Muniz-Rascado, L.; Martinez-Flores, I.; Salgado, H.; Bonavides-Martinez, C.; Abreu-Goodger, C.; Rodriguez-Penagos, C.; Miranda-Rios, J.; Morett, E.; Merino, E.; Huerta, A. M.; Trevino-Quintanilla, L.; Collado-Vides, J., RegulonDB (version 6.0): gene regulation model of Escherichia coli K-12 beyond transcription, active (experimental) annotated promoters and Textpresso navigation. Nucleic Acids Res 2008, 36 (Database issue), D120–4.

24. Cock, P. J.; Antao, T.; Chang, J. T.; Chapman, B. A.; Cox, C. J.; Dalke, A.; Friedberg, I.; Hamelryck, T.; Kauff, F.; Wilczynski, B.; de Hoon, M. J., Biopython: freely available Python tools for computational molecular biology and bioinformatics. Bioinformatics 2009, 25 (11), 1422–3.

